# Melanin-concentrating hormone inhibits PVN oxytocin neurons through a barium-sensitive inwardly rectifying potassium channels and MCH-neuron ablation alters pup-directed aggression

**DOI:** 10.64898/2026.08.15.745000

**Authors:** Tingbi Xiong, Fumihito Saitow, Ayumu Inutsuka, Tatsushi Onaka, Kazuo Yamada, Chitose Orikasa

## Abstract

Interactions between melanin-concentrating hormone (MCH) neurons and oxytocin neurons are crucial for parental care. Whole-cell patch-clamp recordings demonstrated that MCH inhibits paraventricular hypothalamic nucleus (PVN)-oxytocin neurons through activation of barium-sensitive inwardly rectifying potassium channels, potentially G-protein coupled inwardly rectifying potassium channels, and pup-directed aggression was positively related to loss of MCH neurons. Our findings offer a glimpse into the neural mechanisms underlying the evolutionary regulation of offspring caregiving and abuse in males.

## 1. Introduction

Parental behavior, a conserved behavior throughout the animal kingdom, is essential for offspring survival^1^. Sexually inexperienced male mice occasionally exhibit infanticide behavior, however, males with sexual experience and prolonged co-housing with pregnant females gradually undergo a behavioral transition, characterized by a progressive reduction in pup-directed aggression, and is completely abolished following the birth of their offspring^2^. This suppression of aggression can be mimicked by artificial activation of paraventricular hypothalamic nucleus (PVN)-oxytocin neurons in virgin male mice, which is sufficient to inhibit attacking behavior and induce robust parental behavior^3,4^. Accumulating evidence suggests that the interactions between lateral hypothalamic area (LHA)-melanin-concentrating hormone (MCH) neurons and PVN-oxytocin neurons contribute to this marked behavioral shift from aggression to parental behavior. For instance, inputs from LHA-MCH neurons to oxytocin neurons are enhanced after males become fathers^3^. In contrast, knockout MCH neurons induce attacking and increase pup mortality and cannibalism^5,6^. Previous findings have further demonstrated that ablation of MCH neurons is sufficient to abolish PVN-oxytocin neuron mediated parental behavior, which is induced by artificial activation in virgin male mice^4^. Despite these observations, the relationship between LHA-MCH neurons and PVN-oxytocin neurons remains poorly understood. Therefore, we employed electrophysiological approaches to characterize the functional interaction between MCH and PVN-oxytocin neurons and to assess the behavioral consequences of MCH neuron ablation on parental care and aggression in virgin male mice.

## 2. Results

### MCH-induced hyperpolarization in PVN-oxytocin neurons is likely associated with the activation of inwardly rectifying potassium channels

To determine the relationship between MCH and PVN-oxytocin neurons, whole-cell patch-clamp recordings were applied to assess the effect of MCH on the activity of PVN-oxytocin neurons (Fig. 1a). In current-clamp mode, a subset of the recorded PVN-oxytocin neurons responded to MCH with obvious hyperpolarization (9 of 11, 81.8% of recorded neurons), and their mean action potential firing frequency before MCH application was 2.02 ± 0.35 Hz (n = 11). It is speculated that two cells that did not exhibit a hyperpolarization response to MCH were likely TdTomato-negative (oxytocin- nonproducing) neurons that were mistakenly targeted. Given the sparse distribution of TdTomato-positive oxytocin neurons within the PVN, it is highly likely that during patch-clamp recordings, a seal was formed on a non-fluorescent neuron located immediately behind or adjacent to the intended TdTomato-positive target cell. Changes in the membrane potential elicited by 3μM MCH (for 5min), even when non-responders are included, hyperpolarized from -57.5 ± 1.33 mV to -63.2 ± 2.0 mV (t (10) = 3.73, *p* = 0.0039, Fig. 1b, c). In the presence of gabazine (2 μM), which blocks GABAergic synaptic transmission, MCH-induced hyperpolarization persisted. In contrast, application of Ba^2+^ (200 μM), a non-specific GIRK (G-protein coupled inwardly rectifying potassium) channel blocker, reversed the MCH-induced hyperpolarization (4 in Fig. 1b). These results indicate that brief MCH application produces long-lasting inhibitory effects on PVN-oxytocin neurons mediated by GIRK channels rather than GABA_A_ receptors-mediated tonic inhibition. To assess whether MCH directly induces outward currents in PVN-oxytocin neurons, voltage-clamp recordings were conducted to examine MCH-induced membrane currents of PVN-oxytocin neurons. Bath application of MCH (3μM, for 5min) evoked a robust outward current of 161.90 ± 57.83 pA measured at 5 min after onset of MCH application. Subsequent application of Ba^2+^ (200 μM, 5min) rapidly reduced the MCH- evoked outward current to 38.17 ± 17.89 pA measured at 5 min after onset of Ba^2+^ application (n = 6, Fig. 1d). These data also suggest that the sustained outward current induced by MCH is potentially mediated by GIRK channels. To further characterize the ionic mechanism underlying MCH-induced inhibition, current-voltage (I-V) relationships were assessed using voltage-ramp protocols. The reversal potential of MCH-induced currents was -96.6 ± 2.50 mV (n = 10, Fig. 1e, f). These values were close to the predicted equilibrium-potential for K^+^ in our experimental condition (E_K_ = -103 mV). Collectively, these data are consistent with the possibility that MCH activates GIRK channels in PVN-oxytocin neurons, generating outward currents that underlie the observed membrane hyperpolarization and inhibitory effect (Fig. 1g).

**Fig. 1:**
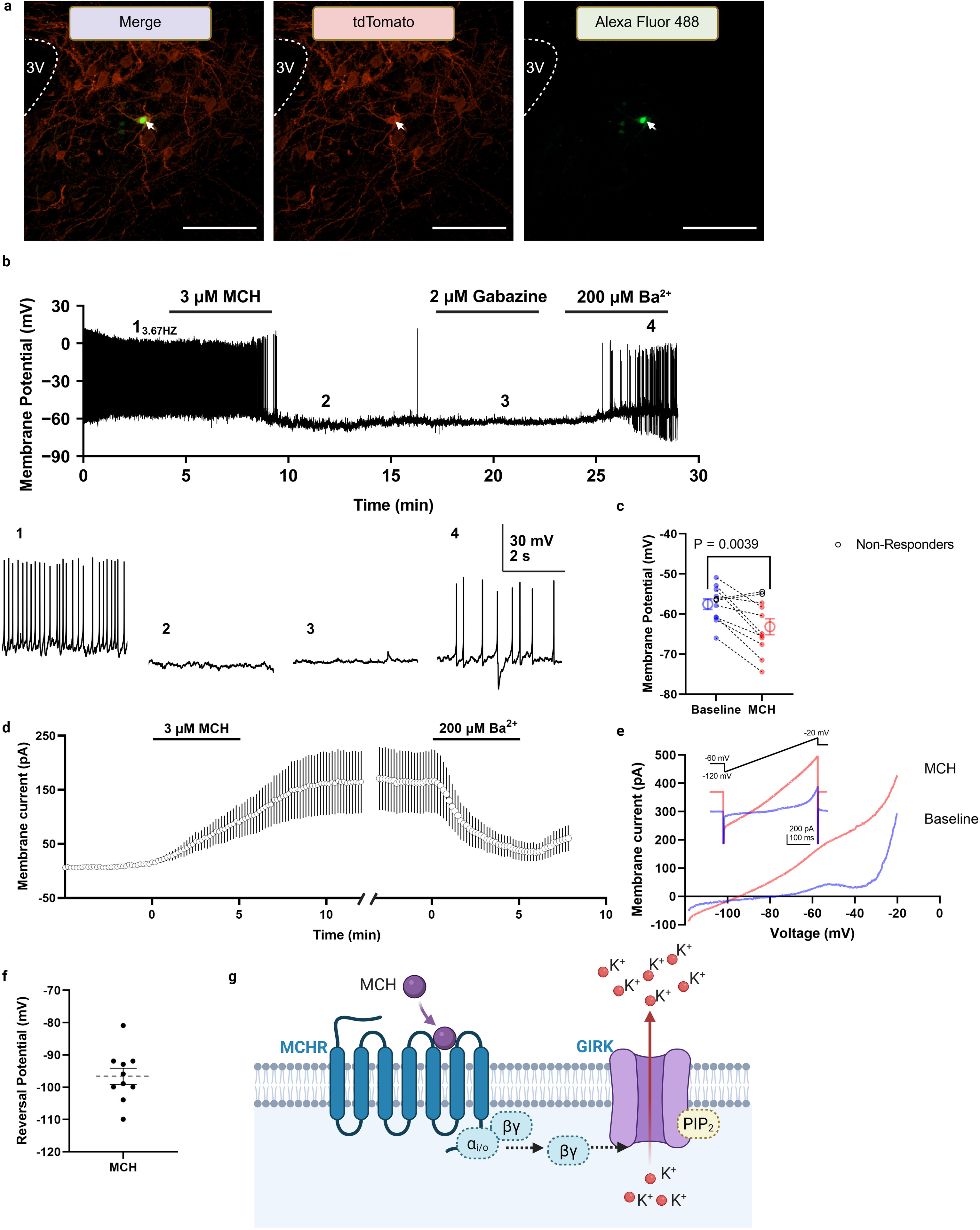
MCH-induced hyperpolarization is reversed by inward rectifier K channel blockade. (**a**) Left panel: merge at 10 × magnification. Red: PVN-tdTomato expressing oxytocin neurons. Green: Alexa Fluor 488 filled into the recorded neurons. Merge: co-localization of tdTomato and Alexa Fluor 488. Arrowheads: simultaneous visualization of PVN-tdTomato expressing oxytocin neurons and PVN-Alexa fluor 488 filled recorded neurons. Middle panel: PVN-tdTomato expressing oxytocin neurons. Right panels: PVN-Alexa fluor 488 filled recorded neurons. Scale bars, 100 μm. (**b**) Upper panel: representative trace showing that application of MCH (3 μM) induced membrane hyperpolarization and suppressed action potential firing in PVN-oxytocin neurons (lower panels 1 and 2). The MCH-induced hyperpolarization persisted in the presence of gabazine (2 μM; lower panel 3). Subsequent application of Ba^2+^ (200 μM) reversed the MCH-induced hyperpolarization (lower panel 4). Lower panel 1: baseline depolarized membrane potential of a PVN-oxytocin neurons. Lower panel 2: hyperpolarization induced by MCH. Lower panel 3: blockade of GABAergic synaptic transmission didn’t affect MCH-induced hyperpolarization. Lower panel 4: blockade of an inward rectifier K^+^ channels reversed the MCH-induced hyperpolarization. (**c**) Extent of MCH-induced hyperpolarization in PVN-oxytocin neurons. Paired t test, t(10) = 3.73, *p* = 0.0039, n = 11 (Responder n = 9, Non-responder n = 2). All data are presented as means ± SEM. (**d**) Application of MCH (3 μM) induced a persistent outward current, which was rapidly reduced by subsequent application of Ba^2+^ (200 μM). (**e**) Current recorded by a voltage ramp (400 ms) from -120 mV to -60 mV before (blue line: Baseline) and during (red line: MCH) MCH application in a PVN-oxytocin neuron. Current-voltage relationship showing the increase in membrane conductance induced by MCH (3 μM). (**f**) The reversal potential of MCH evoked current was -96.6 ± 2.50 mV. All data are presented as means ± SEM. n = 10. (**g**) Schematic illustration of the proposed mechanism of MCHR1-mediated signaling via inwardly rectifying potassium channels, potentially GIRK channels, in PVN-oxytocin neurons. All data are presented as means ± SEM.

### Ablation of MCH neurons induced aggression

In the behavior assays (Fig. 2a, b), no significant differences were observed among groups with partial or complete ablation of MCH neurons and the control group in positive parental behaviors, including retrieving and crouching, or negative burying behavior (Fig. 2c, d, e). However, complete ablation of MCH neurons resulted in a significant increase in pup-directed attacking compared with the control group. These results were confirmed by a Chi-square test followed by Fisher’s exact *post hoc* test (χ^2^(2) = 6.233, *p* = 0.0443; Fig. 2f). In partial and complete ablation groups, the number of MCH neurons significantly reduced compared with control groups, and this result was confirmed by one-way ANOVA followed by Bonferroni’s *post hoc* tests (F(2, 20) = 94.90, *p* < 0.0001; Fig. 2g).

**Fig. 2:**
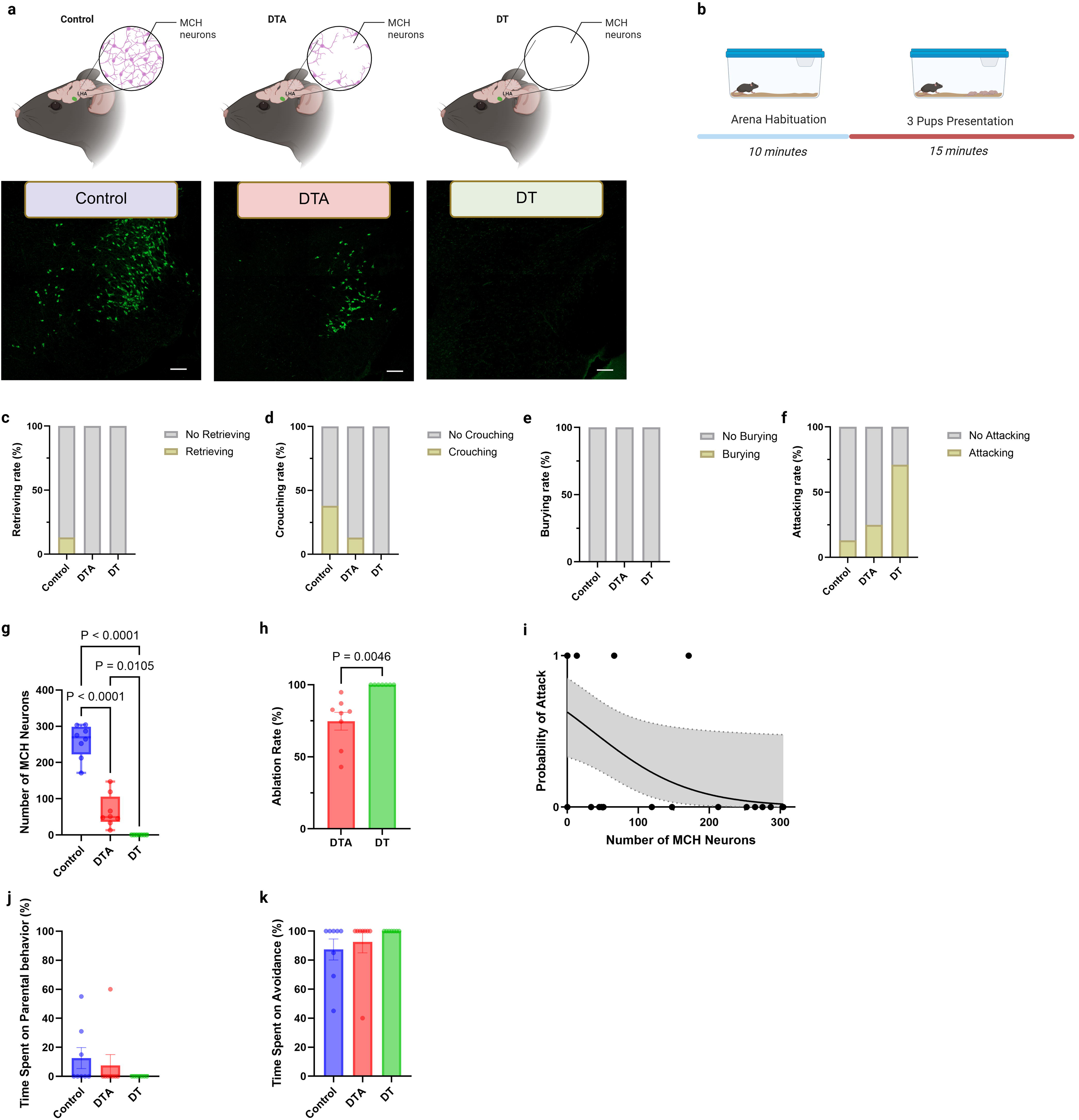
Ablation of MCH neurons induced aggression. (**a**) Green signal indicating the LHA-MCH- ir neurons following different treatments. Left panel: control group without MCH neurons ablation (n = 8). Middle panel: partial ablation group (DTA; n = 8). Right panel: complete ablation group (DT; n = 7). Scale bars, 100 μm. (**b**) Graphic representation of parental behavior test. (**c**) Graphic displaying the rate of retrieving behavior after different treatments. Chi-square test, χ^2^ = 1.960, df = 2, ns. (**d**) Graphic showing the rate of crouching following different treatments. Chi-square test, χ^2^ = 3.859, df = 2, ns. (**e**) Graphic showing the rate of burying after different treatments. (**f**) Graphic exhibiting the rate of attacking after different treatments. Chi-square test, χ^2^(2) = 6.233, *p* = 0.0443. (**g**) Graphic showing the quantification of MCH-ir neurons after different treatments. One- way ANOVA, F(2, 20) = 94.90, *p* < 0.0001 (**h**) Graphic representation of the ablation ratio of LHA– MCH neurons in different groups. Unpaired t test, t(7) = 4.102, *p* = 0.0046 (**i**) Graphic showing of the relationship between the number of MCH neurons and the probability of attack. Univariable logistic regression, *G*^2^ = 7.51, *p* = 0.0062 (**j**) Graphic exhibiting of the percentage parental behavior after different treatments. One-way ANOVA, F(2, 20) = 0.9828, ns. (**k**) Graphic showing of the percentage avoidance after different treatments. One-way ANOVA, F(2, 20) = 0.9828, ns. All data are presented as means ± SEM.

The ablation rate of MCH neurons in partial and complete ablation group was approximately 75% and 100%, respectively (t(7) = 4.102, *p* = 0.0046; Fig. 2h). Because pup-directed attacking was changed among groups, the relationship between the number of MCH neurons and the probability of attacking was further examined. An univariable logistic regression analysis revealed a significant negative association between the number of MCH neurons and attacking behavior (*G*^2^ = 7.51, *p* = 0.0062; Fig. 2i). Each additional MCH neurons was associated with an odds ratio of 0.9855 for attacking (95% CI, 0.9658-0.9966), corresponding to an approximately 1.45% decrease in the odds of attacking per additional neuron. The regression equation was log[p/(1 - p)] = 0.5231 – 0.01459X, where p presents the probability of attacking and X represents the number of MCH neurons. No significant difference was detected in sniffing latency and sniffing duration among these groups (Supplementary Fig. S1a, b). Heat map analyses of spatial occupancy revealed that the control group spent a modulate amount of time near the pups, whereas mice with partial ablation of MCH neurons exhibited avoidance behavior and spent more time in the area of the cage which is opposite to the pups. In contrast, mice with complete ablation of MCH neurons continuously moved along the periphery of the cage and displayed hyperactivity (Supplementary Fig. S2a). Locomotor analysis showed a significant increase in movement speed in the complete ablation group compared with the partial ablation and control groups. These results were determined by one-way ANOVA followed by Bonferroni’s *post hoc* tests (*F* (2, 20) = 10.44, *p* <0.001; Supplementary Fig. S2b). No significant difference was detected in duration of close to pup among these groups (Supplementary Fig. S2c).

We defined the average time spent on burying and moving away from the pups as “Avoidance” and the time spent on retrieving and crouching as “Parental behavior” and then calculated the ratio of these behaviors. No significant difference was detected in time spent on parental behavior and avoidance (Fig. 2j, k).

### Ablation of MCH neurons did not affect adult-directed aggression

In addition, to determine whether complete ablation of MCH neurons also enhances aggression toward adult conspecifics, a resident-intruder assay was performed. (Fig. 3a, b). No significant differences were observed in sniffing latency or duration between groups (Supplementary Fig. S3a, b). And no significant differences were observed in either severe or mild aggression between groups (Supplementary Fig. S4, 5; Supplementary Table S1). Together, these results suggest that complete ablation of MCH neurons didn’t affect adult-directed aggression.

**Fig. 3:**
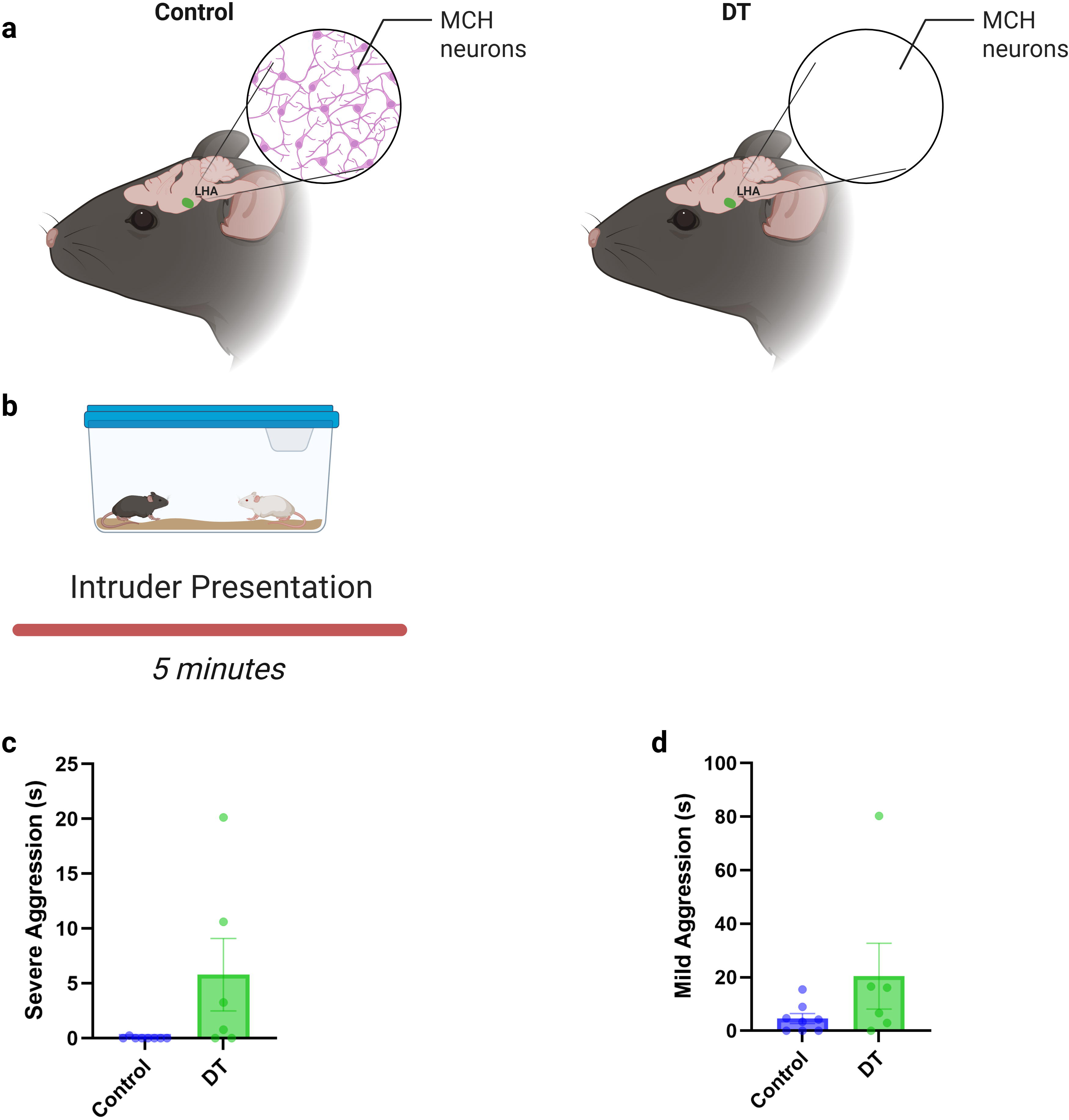
Ablation of MCH neurons induced adult-directed aggression tendency. (**a**) Left panel: control group without MCH neurons ablation (n = 8). Right panel: complete ablation group (DT; n = 6). (**b**) Schematic representation of resident-intruder assay. (**c**) Graphs representing the duration of severe aggression after different treatments. Unpaired t test, t(12) = 2.044, ns. (**d**) Graphs exhibiting the duration of mild aggression after different treatments. Unpaired t test, t(12) = 1.478, ns. All data are presented as means ± SEM.

## 3. Discussion

In the present study, bath application of MCH stably suppressed PVN-oxytocin neuron excitability. Voltage-clamp conditions further revealed that MCH activated outward currents that reversed near the potassium equilibrium potential and were reduced by Ba^2+^, but not by gabazine. These findings suggest that MCH released by hypothalamic neurons inhibits PVN-oxytocin neurons via enhanced activity of Ba^2+^-sensitive inwardly rectifying potassium channels, potentially of the GIRK channels. Consistent with previous reports^4,6^, parental behavior was impaired following partial or complete cell-type specific ablation of MCH neurons. And ablation of MCH neurons promoted pup-directed aggression depends on the extent of MCH neuron loss.

MCH receptor 1 (MCHR1) is a seven-transmembrane domain G-protein-coupled receptor^7^. Binding of MCH to MCHR1 activates Gi-, Go-, and Gq-dependent intracellular signaling pathways, which mediate diverse physiological functions^8^. Payant et al. (2024) reported that inhibitory actions of MCH in the lateral septum are mediated by protein kinase C-dependent Gq coupling and subsequent activation of GABA_A_ receptors^9^. Previous studies have shown that MCH neurons contain GABA^10^, and that their terminals in the PVN are GABAergic^6^, suggesting that MCH-induced inhibition of PVN-oxytocin neurons might be through GABAergic synaptic transmission. However, application of gabazine did not alter MCH-induced hyperpolarization in PVN-oxytocin neurons, indicating that alternative mechanisms are involved. Prior evidence has suggested that the inhibitory effects of MCH neurons may also be mediated through potassium ion efflux^11^. Consistent with this possibility, application of Ba^2+^ reversed MCH-induced hyperpolarization, suggesting the involvement of inwardly rectifying potassium channels, potentially GIRK channels. However, it should be noted that Ba^2+^ are broad inhibitors of the inwardly rectifying potassium channel family; thus, the specific contribution of GIRK channels remains a likely possibility rather than a definitive conclusion due to these pharmacological limitations. Study shows that activation of GIRK channels typically requires phosphatidylinositol 4,5-bisphosphate (PIP2) and occurs through the biding G-protein βγ subunits following activation of G-protein-coupled receptors^12^. The observation of MCH-induced putative GIRK currents in oxytocin neurons is further supported by previous findings that MCH binding to MCH receptors induces potassium channel-mediated currents^13^, suggesting coupling of MCHR1 to Gi/o-dependent signaling pathway in PVN-oxytocin neurons. Accumulating evidence indicates that reduced GIRK signaling promotes neuronal hyperexcitability and facilitates aversive encoding^14,15^. Therefore, ablation of MCH neurons may diminish GIRK-mediated inhibitory signaling and promoting hyperexcitability in PVN-oxytocin neurons.

Initiation of normal parental behavior requires coordinated activity between oxytocin and MCH neurons. Virgin rodents of both sexes typically avoid pups rather than giving parental care^16,17^. Consistent with this, sexually inexperienced male mice were found to exhibit limited parental behaviors toward pups. When LHA-MCH neurons are intact, artificial activation of oxytocin neurons enables virgin mice of both sexes to express robust parental behavior toward pups^4,18^. However, even partial ablation is sufficient to disrupt oxytocin neurons induced parental behavior^4^. Previous studies have demonstrated that the MCHR1 but not MCH receptor 2 (MCHR2) is widely expressed in rodents^19^, and knockout of this MCHR1 was reported to impair the parental behavior of both postpartum and virgin female mice^20, 21^. The absence of detectable differences in parental behavior among control, partial, and complete ablation groups in virgin male mice is likely attributable to a floor effect, as virgin males normally exhibit very limited parental behavior compared with oxytocin neuron activated animals or parents with reproductive experiences.

Although pup vocalizations could promote parental behavior through sustained disinhibition of oxytocin neurons^22^, and many studies have reported an inverse relationship between oxytocin neurons activity and aggression, with higher oxytocin signaling associated with reduced aggression^23,24^. Because MCH inhibits PVN-oxytocin neurons, theoretically, ablation of MCH neurons will disinhibit oxytocin neurons, which in turn promote social behavior and inhibit aggression. However, instead of promoting social behavior, the effects of MCH neuron ablation in the present study show pup-directed aggression were obvious. To deal with this discrepancy, we propose that the increased aggression demonstrated by mice with complete ablation of MCH neurons may result from loss of critical inhibitory input to PVN-oxytocin neurons and possibly leads to hyperactivation. That is, slightly removed inputs from MCH neurons may result in modulate excitation on PVN-oxytocin neurons, which facilitate pro-social behavior. However, increasing the ablation rate will further disinhibition of oxytocin neurons and eventually results in activation of oxytocin neurons exceed appropriate range for occurring pro-social behavior, and results in negative impact. This inverted U-shape relation between oxytocin neuron activation and social behavior may be the explanation of the discrepancy. In fact, the emerging evidence indicates that activation of oxytocin neurons can facilitate aggression. For instance, Sofer et al. (2024) reported that activation of PVN-oxytocin neurons in wild house mice is sufficient to induce aggressive behavior toward both pups and other adults^25^. In addition, Bosch et al. (2005) reported that heightened aggression in rats with high anxiety-related behavior is accompanied by increased oxytocin release into the extracellular space of the PVN^26^.

MCH neurons robustly express oxytocin receptors and can be directly excited by oxytocin neurons through synaptic activation of Na^+^/Ca^2+^ exchangers and nonselective cation channels^27, 28, 29^. Hypothalamic PVN-oxytocin neurons express MCHR1 and receive direct projections from MCH neurons^6,30^. These findings combined with present electrophysiology experiment results that PVN- oxytocin neurons are directly inhibited by MCH via presumably GIRK channels but not through GABAergic transmissions, may supporting the existence of an oxytocin-MCH feedback mechanism in the regulation of pup-directed behavior. Such a reciprocal system may reconcile long-lasting contradictions regarding the functions of oxytocin in parental and aggressive behaviors.

We suggest that pup vocalizations likely activate oxytocin neurons, which in turn stimulate MCH neurons. Increased activation of MCH neurons then provides inhibitory feedback to modulate oxytocin neuron excitability within an appropriate range, thereby suppressing aggression and promoting parental behavior. However, the complete loss of MCH neurons prevents this inhibitory modulation of oxytocin neurons, resulting in hyperexcitability and aggression toward pups. Alternatively, partial ablation of MCH neurons may preserve limited feedback control, thereby preventing overt aggression.

In addition, complete ablation increased locomotor activity is consistent with prior studies in MCH knockout mice^31, 32^. Loss of dopaminergic innervation is well known to produce motor deficits in Parkinson’s disease^33^, whereas dopamine exerts pro-locomotor effects primarily through activation of D1 receptors^34, 35^. Activation of oxytocin neurons terminals in the ventral tegmental area (VTA) is sufficient to enhance dopamine neuron activity^36^ and to induce aggressive behavior^25^. Moreover, knockout of MCH has been shown to evoke robust dopamine release^37^. The elevated locomotor activity and aggression observed after MCH neuron ablation may therefore be explained, at least in part, by the proposed oxytocin-MCH feedback mechanism, in which loss of MCH-mediated inhibition results in hyperexcitability of PVN-oxytocin neurons.

In summary, the present study suggested that PVN-oxytocin neurons are directly inhibited by MCH via enhanced activity of Ba^2+^-sensitive inwardly rectifying potassium channels, with GIRK channels being the most likely candidates. MCH neuron ablation impaired parental behavior and promoted aggression toward pups. Future studies may employ in vivo calcium imaging combined with virus tracing approaches. This may help confirm that MCH deficiency-induced aggression arises from the hyperexcitability of PVN-oxytocin neurons and downstream targets such as VTA or the bed nucleus of the stria terminalis^25, 38^. Selected GIRK channel blocker should also be applied to determine whether MCH-mediated inhibition of oxytocin neurons depends on GIRK channel activation. In addition, projection-specific approaches could be used to examine whether pup-directed behaviors are altered by the selective loss of MCH input to PVN-oxytocin neurons. Our findings offer a glimpse into complex interactions between oxytocin and MCH neurons in regulating social behavior and provide a conceptual framework for reconciling the observed aggressive effects of MCH neurons.

## 4. Materials and Methods

### Subjects

All housing and experimental protocols were approved by the Nippon Medical School Committee on Animal Research (Approval No. 2026-011) and were conducted in compliance with the Guidelines for the Care and Use of Laboratory Animals of the Japanese Ministry of the Environment (Notification No. 88, 2006) and related regulations. This study is reported in accordance with the ARRIVE guidelines. Sexually naïve male mice were housed in 19 × 27 × 15 cm transparent polypropylene cages under controlled temperature (23°C) and humidity (50 ± 10%) on a 12-h light/dark cycle (lights on at 23:00), with food and water available *ad libitum*. All experiments were conducted during the dark phase. The inbred strains of mice were bred in our laboratory.

### Transgenic lines

To specifically target PVN-oxytocin neurons for electrophysiology recordings, Oxytocin-ires-Cre mice were used [B6;129S-Oxttm1.1(cre)Dolsn/J, 024234] (Jackson Laboratory, Ellsworth, ME, USA). For partial ablation of LHA-MCH neurons, previously established double transgenic omb-Cre mice^4^, which express Cre recombinase under control of the oxytocin and MCH promoters, were used [MCH-Cre, STOCK Tg(Pmch-cre)1Lowl/J, 014099] (Jackson Laboratory, Ellsworth, ME, USA). For complete congenital ablation of LHA-MCH neurons, previously established DT mice were used; these mice eliminated MCH neurons via a Tet-off system generated by breeding MCH-tTA mice (gifted from Dr. Yamanaka), with TetO-DTA [B6.Cg-Tg(tetO-DTA)1Gfi/J, 008468] (Jackson Laboratory, Ellsworth, ME, USA)^6^.

### Type of viruses

The adeno-associated virus (AAV) Helper-Free system (Agilent Technologies, Inc., Santa Clara, CA, USA) was used to produce and purify AAV vectors. AAV9-CAG-FLEX-tdTomato-CAAX-WPRE and AAV9-CMV-FLEX-hrgreen fluorescent protein (GFP) were used to express red fluorescent protein or hrgreen fluorescent protein in a Cre recombinase-dependent manner, respectively. AAV9- CMV-FLEX-DTA was applied to ablate neurons in a Cre recombinase-dependent manner. The efficacy of the DTA for neuron-specific ablation was confirmed as previously reported^4, 6, 39^.

### Stereotaxic AAV injection

All injections were performed using a stereotactic frame (Devid Kopf Instruments, Tujunga, CA, USA, with Stoelting Mouse and Neonatal Rat Adaptor, Wood Dale, IL, USA), 27-gauge stainless steel tubing, and a microprocessor-controlled syringe pump (World Precision Instruments, Inc., Sarasota, FL, USA) were used at a rate of 0.12 μL/min. AAV9-CAG-FLEX-tdTomato-CAAX- WPRE treated animals were injected at 5 weeks of age and used for electrophysiology recordings at 9 weeks. AAV9-CMV-FLEX-hrGFP or AAV9-CMV-FLEX-DTA treated animals were injected at 8 weeks of age and were tested for behavior at 12-16 weeks.

To express tdTomato in PVN-oxytocin neurons, Oxytocin-ires-Cre mice were bilaterally injected with AAV9-CAG-FLEX-tdTomato-CAAX-WPRE (1.1 × 10^13^ copies/mL) into PVN (relative to bregma: anteroposterior, − 0.6 mm; mediolateral, ± 0.2 mm; dorsoventral, − 4.6 mm) at 600 nL per injection.

For selective ablation of LHA-MCH neurons, omb-Cre mice were bilaterally injected with AAV9- CMV-FLEX-DTA (5.3 × 10^12^ copies/mL) or AAV9-CMV-FLEX-hrGFP (5.3 × 10^12^ copies/mL) into the LHA (relative to bregma: anteroposterior, − 1.5 mm; mediolateral, ± 0.9 mm; dorsoventral, − 5.0 mm) at 600nL per injection.

### Electrophysiology in the PVN

Male mice at 9 weeks of age were deeply anaesthetized by isoflurane inhalation (approximately 4% in air, v/v) and closely monitored until loss of the limb withdrawal reflex following toe pinch was confirmed. Then, they were transcardially perfused with 10 mL of ice-cold oxygenated NMDG (N-Methyl-D-glucamine) cutting solution^40^ that contained the following compounds: 110 mM NMDG, 30 mM NaHCO_3_, 2.5 mM KCl, 0.5 mM CaCl_2_, 10.0 mM MgCl_2_, 1.25 mM NaH_2_PO_4_, 12.5 mM glucose and 20 mM HEPES (PH 7.4). Brains were rapidly removed from the skull and sliced coronally at 280 μm thickness using a vibratome (VT1200S; Leica Biosystems, Nussloch, Germany) at 4°C in the NMDG cutting solution. Slices containing the PVN were transferred to a submerged chamber for at least 1.5 h in artificial cerebrospinal fluid (ACSF) that contained 125.0 mM NaCl, 2.5 mM KCl, 2 mM CaCl_2_, 1.3 mM MgCl_2_, 26.0 mM NaHCO_3_, 1.25 mM NaH_2_PO_4_, and 11.0 mM glucose. ACSF was maintained at pH 7.4 by bubbling 95% O_2_-5% CO_2_ gas. Except for the membrane potential measurements using the current-clamp mode (Fig. 1b, c), slices were superfused with ACSF containing 500 nM tetrodotoxin (FUJIFILM Wako Pure Chemical, Tokyo, Japan) to block voltage-gated Na^+^ channels and eliminate action potential-dependent synaptic activity.

Slices containing the PVN were bisected along the midline and transferred to the recording chamber mounted on the stage of a microscope (40×, NA = 0.80; Olympus, Tokyo, Japan). Slices were continuously perfused with ACSF at a flow rate of 1.4 ml/min at 30 °C. Whole-cell patch-clamp recordings were obtained with borosilicate glass pipettes (3-5 MΩ; World Precision Instruments, Sarasota, USA) filled with a potassium-based internal solution (containing: 150 mM potassium methanesulfonate, 5.0 mM KCl, 0.1mM K-EGTA, 5.0 mM Na-HEPES, 3.0 mM Mg-ATP, 0.4 mM Na-GTP, pH 7.4). Cells that were visualized under green light were considered tdTomato-positive and were designated as oxytocin neurons. During the recordings, recorded neurons were infused with 15 μM Alexa Fluor 488 (Thermo Fisher Scientific, Waltham, USA) to determine whether the neurons were oxytocin neurons (Fig. 1a). Whole-cell patch-clamp recordings were acquired and controlled using the Axon 700B Multiclamp amplifier (Molecular Devices, San Jose, USA) and pClamp10 acquisition software (Molecular Devices). The membrane potentials were measured in the current- clamp mode. The effects of MCH receptor activation on the membrane potential were determined by recording from 11 neurons in three animals (Fig. 1b-1c). Neurons were classified as responsive if MCH induced a sustained hyperpolarization and decreasing the firing frequency of action potentials. For recordings of MCH-induced currents in the voltage-clamp mode, the cells were voltage clamped at −59.9 mV. Data were obtained from 6 cells derived from two animals (Fig. 1d-1f). Data acquisition and analysis were performed using Clampfit (Molecular Devices) and KyPlot (KyensLab Inc., Tokyo, Japan). All signals were filtered at 2 kHz and sampled at 10 kHz. The membrane potential was shown as a value corrected for the liquid junction potential. In our previous measurement, it was +9.9 mV^41^.

After establishment of a stable baseline, MCH (3 µM, MedChemExpress, Monmouth Junction, NJ, USA) was bath applied into the recording chamber for 5 min, followed by a washout period with ACSF. Gabazine (2 µM, Sigma-Aldrich, St. Louis, MO, USA) and Ba^2+^ (200 µM, FUJIFILM Wako Pure Chemical, Tokyo, Japan) were subsequently applied to the slice for 5 min following MCH application.

### Standard behavioral assays

#### Parental behavior

A cage containing the mouse was transported from the shelves to the observation area and was allowed to habituate for 10 min to the experimental setting. The test started by placing three 3- to 5- day-old pups on the opposite side of the mouse for the next 15 min. The distance recorded between the mouse and the pups was defined as “close” when staying within proximity (< 1 cm) of pups.

Parental behavior was assessed over 15 min by observing retrieving and crouching behaviors as previously reported^42^. Virgin male mice often exhibited limited parental behaviors toward pups compared with sires. Parental behavior was categorized according to the following criteria: test mice that picking up a pup with their mouth and carrying it to the nest were categorized as “retrieving”, and animals that crouched over pups with their limb extension and assuming a nursing-like posture were categorized as “crouching”. Ignore behavior was defined as remaining more than 1 cm away from pups and exhibiting no response to pups. Burying behavior was defined as the mouse directing its head toward a pup and using its front limbs to scoop bedding toward the pup until it is completely covered by the bedding and no longer visible. Attacking behavior was defined as using the teeth to seize and clamp down on the pup. Attacked pups were immediately removed from the test cage and the tests were ended. Licking behavior was defined as sniffing or licking.

#### Resident-intruder assay

As in the procedure above, the cage containing the mouse was transferred to the observation area. An unfamiliar mouse that had undergone olfactory bulbectomy was then introduced into the home cage for 5 min. Aggressive behaviors including lateral threat, chase, tug, launch, bite, and clinch, were recorded as previously reported^25^. Lateral threat was defined as assuming a lateral posture and the threatened mouse stands on its hind legs in a defensive posture. Chase was defined as a rapid pursuit toward the intruder with the intent to attack, accompanied by the intruder mouse fleeing from the resident. Tug was defined as pulling and dragging the intruder down. Launch was defined as a rapid movement toward the intruder with intent to attack. Bite was defined as seizing and clamping down the intruder with the teeth. Clinch was defined as a rapid attack involving mice grabbing onto each other using forelimbs and rolling in a tight hold. Aggressive behaviors were further categorized as severe aggression (behaviors likely to cause injury, such as bite and clinch) or mild aggression (the remaining aggressive behaviors). Sniffing behavior was also recorded.

### Immunohistochemistry

Serial coronal sections (40 μm) were cut using a Microm HM 560 cryostat (Thermo Fisher Scientific, Waltham, MA, USA). All immunostaining steps were conducted at room temperature and on a plate shaker. Sections were first rinsed three times (10 min per wash) in 0.01M PBS (pH 7.4). For MCH ablation confirmation, sections were incubated overnight with rabbit anti-MCH antibody (1:10,000; Phoenix Pharmaceuticals Inc., Burlingame, CA, USA). On the following day, sections were rinsed three times in PBS and incubated with Alexa Fluor 488-goat anti-rabbit IgG (1:200; Molecular Probes Inc., Eugene, OR, USA). After a final rinse in PBS, stained sections were then wet mounted on slides. Once dried, they were covered with mounting medium (Vector laboratories, inc., Newark, CA, USA).

### Machine Learning

Body-part tracking was performed in DeepLabCut (v2.3.9) with a ResNet-50 network trained for 100,000 iterations on 360 frames labeled at eight landmarks (left ear, right ear, nose, middle body, left body, right body, tail base, tail tip). Training and test errors were 3.26 pixels and 8.61 pixels, respectively. Spatial occupancy, speed, and closeness to pup were analyzed by a custom Python (version 3.8.19) script.

### Statistical Analysis

All data were analyzed using GraphPad Prism (version 10.6.1; GraphPad Software Inc., San Diego, CA, USA). For electrophysiological experiments, membrane potential measured before and during MCH application were compared using paired t test. For the parental behavior experiments, categorical variables were compared between groups by Chi-squared test followed by Fisher’s exact test for *post hoc* analysis. Continuous variables such as the number of MCH neurons, time spent on parental behavior, time spent on avoidance, sniffing latency, sniffing duration, speed, close to pup were compared using one-way analysis of variance (ANOVA), followed by Bonferroni’s *post hoc* tests. MCH neuron ablation rates were compared using an unpaired t test. The relationship between number of MCH neurons and probability of attack was evaluated using univariable logistic regression. For the resident-intruder assay, the duration of severe and mild aggression was compared between groups using unpaired t test. Subcategory of severe and mild behavioral duration and numbers were analyzed using Mann-Whitney U test. Behavior latencies were using Kaplan-Meier time-to-event and compared between groups using two-sided log-rank tests. The Holm-Sidak correction was applied to adjust for multiple comparisons among subcategory behaviors.

## Supporting information

Supplementary Table S1. Statistics

## 5. Data availability

The datasets used and/or analyzed during the current study available from the corresponding author on reasonable request.

## 7. Acknowledgments

This work was partially supported by Division of Advanced Research Promotion, Aichi Medical University.

## 8. Funding

This work was supported, in part, by grants-in-aid for scientific research from the Japanese Ministry of Education, Science, Sports and Culture [23K05858 (2023–2026)] (C.O.).

---

**Supplementary Fig. S1: Pup-directed sniffing behavior was not affected by ablation of MCH neurons.** (**a**) Graphs representing the duration of sniffing latency after different treatments. One-way ANOVA, F(2, 20) = 3.101, ns. (**b**) Graphic showing the duration of sniffing after different treatments. One-way ANOVA, F(2, 20) = 2.524, ns. All data are presented as means ± SEM.

**Supplementary Fig. S2: Locomotor activity was increased by ablation of MCH neurons.** (**a**) Graphs representing spatial occupancy maps after different treatments. The black dots indicate pups’ location (**b**) Graphic showing the moving speed after different treatments. One-way ANOVA, F(2, 20) = 10.44, *p* = 0.0008 (**c**) Graphs exhibiting the duration of close to pups after different treatments. One-way ANOVA, F(2, 20) = 3.438, ns. All data are presented as means ± SEM.

**Supplementary Fig. S3: Adult-directed sniffing behavior was not affected by ablation of MCH neurons.** (**a**) Graphs representing the duration of sniffing latency after different treatments. Unpaired t test, t(12) = 0.4148, ns. (**b**) Graphic showing the duration of sniffing duration after different treatments. Unpaired t test, t(12) = 0.2495, ns. All data are presented as means ± SEM.

**Supplementary Fig. S4: Adult-directed severe aggression was not affected by ablation of MCH neurons.** Graphs representing the bite duration (Mann-Whitney U test, U = 9, ns) (**a**), number of bite (Mann-Whitney U test, U = 9, ns) (**b**), and bite latency (log-rank test, χ²(1) = 5.336, ns) (**c**) after different treatments. Graphic showing the clinch duration (Mann-Whitney U test, U = 16, ns) (**d**), number of clinch (Mann-Whitney U test, U = 16, ns) (**e**), and clinch latency (log-rank test, χ²(1) = 2.925, ns) (**f**) after different treatments. All data are presented as means, and the whiskers indicate the interquartile range.

**Supplementary Fig. S5: Adult-directed mild aggression was not affected by ablation of MCH neurons.** Graphs representing the launch duration (Mann-Whitney U test, U = 16, ns) (**a**), number of launch (Mann-Whitney U test, U = 16, ns) (**b**), and launch latency (log-rank test, χ²(1) = 2.925, ns) (**c**) after different treatments. Graphic showing the threat duration (Mann-Whitney U test, U = 22.5, ns) (**d**), number of threat (Mann-Whitney U test, U = 24, ns) (**e**), and threat latency (log-rank test, χ²(1) = 0.2486, ns) (**f**) after different treatments. Graphic displaying the tug duration (Mann-Whitney U test, U = 20, ns) (**g**), number of tug (Mann-Whitney U test, U = 20, ns) (**h**), and tug latency (log- rank test, χ²(1) = 1.167, ns) (**i**) after different treatments. Graphic exhibiting the chase duration (Mann-Whitney U test, U = 10, ns) (**j**), number of chase (Mann-Whitney U test, U = 9, ns) (**k**), and chase latency (log-rank test, χ²(1) = 4.905, ns) (**l**) after different treatments. All data are presented as means, and the whiskers indicate the interquartile range.

## Notes

### Competing Interest Statement

The authors have declared no competing interest.

### Summary of Updates

This version of the manuscript has been updated to add another supplemental file.

