## Supplementary figures and images for "Melanin-concentrating hormone inhibits PVN oxytocin neurons through a barium-sensitive inwardly rectifying potassium channels and MCH-neuron ablation alters pup-directed aggression"

### Supplemenatry 4

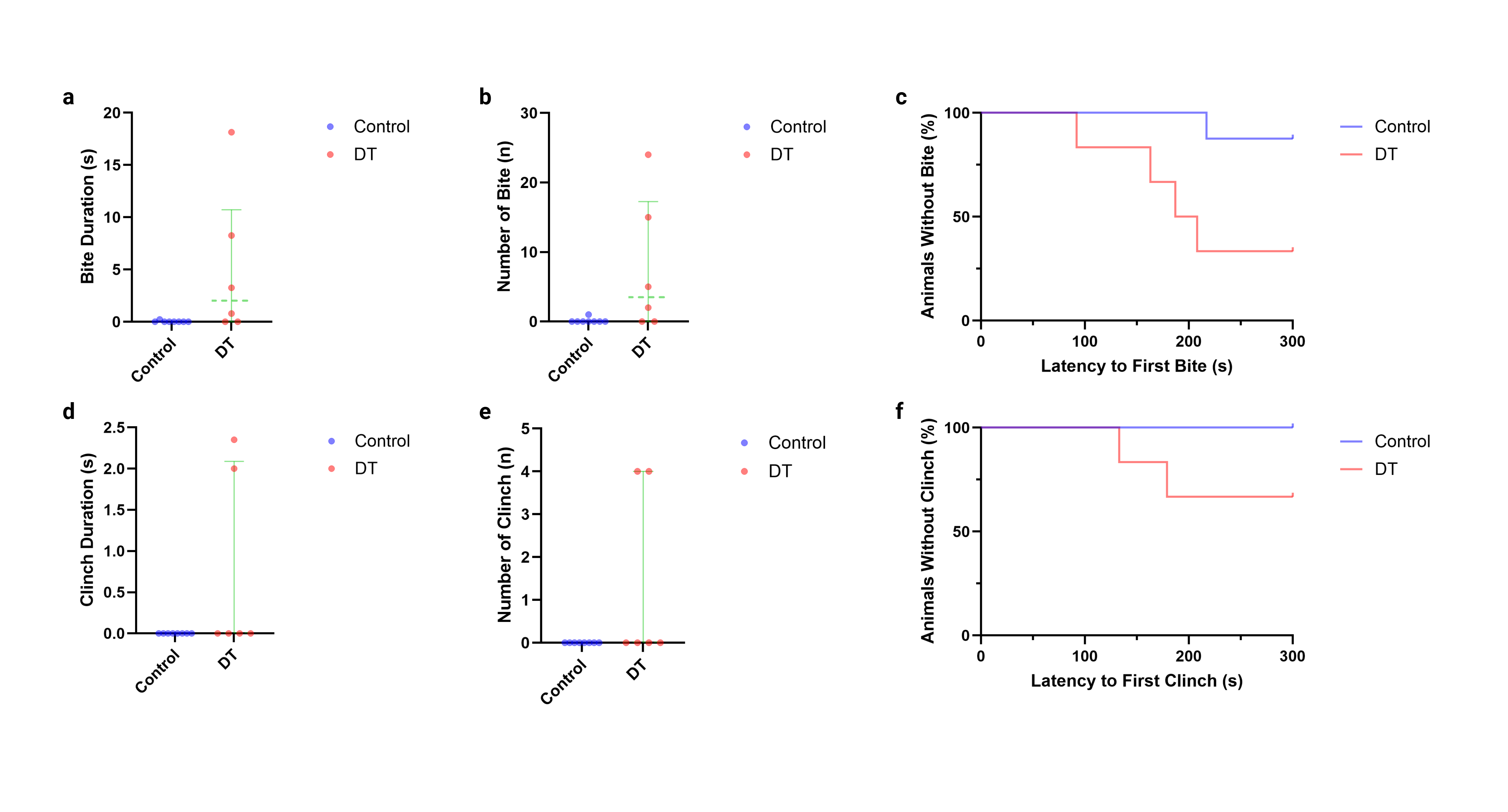

### Supplementary 1

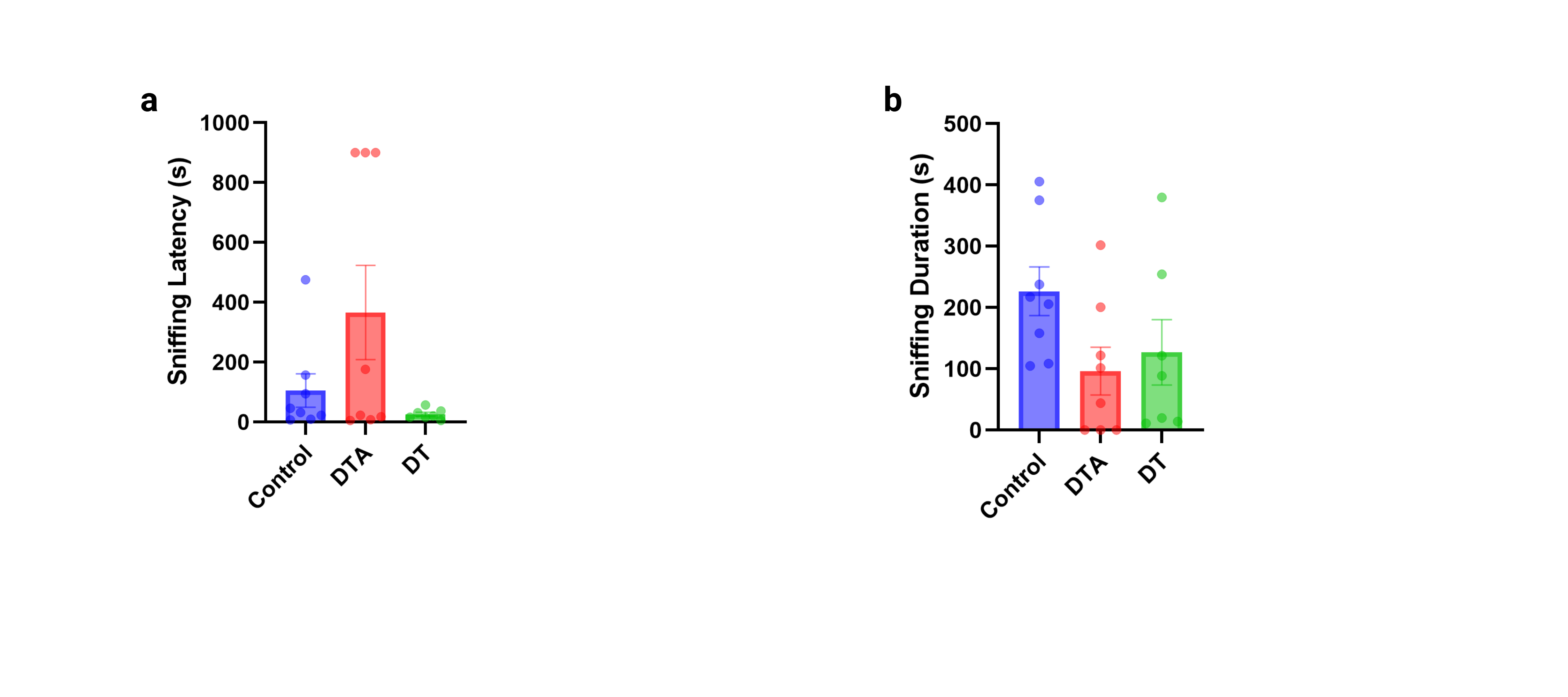

### Supplementary 2

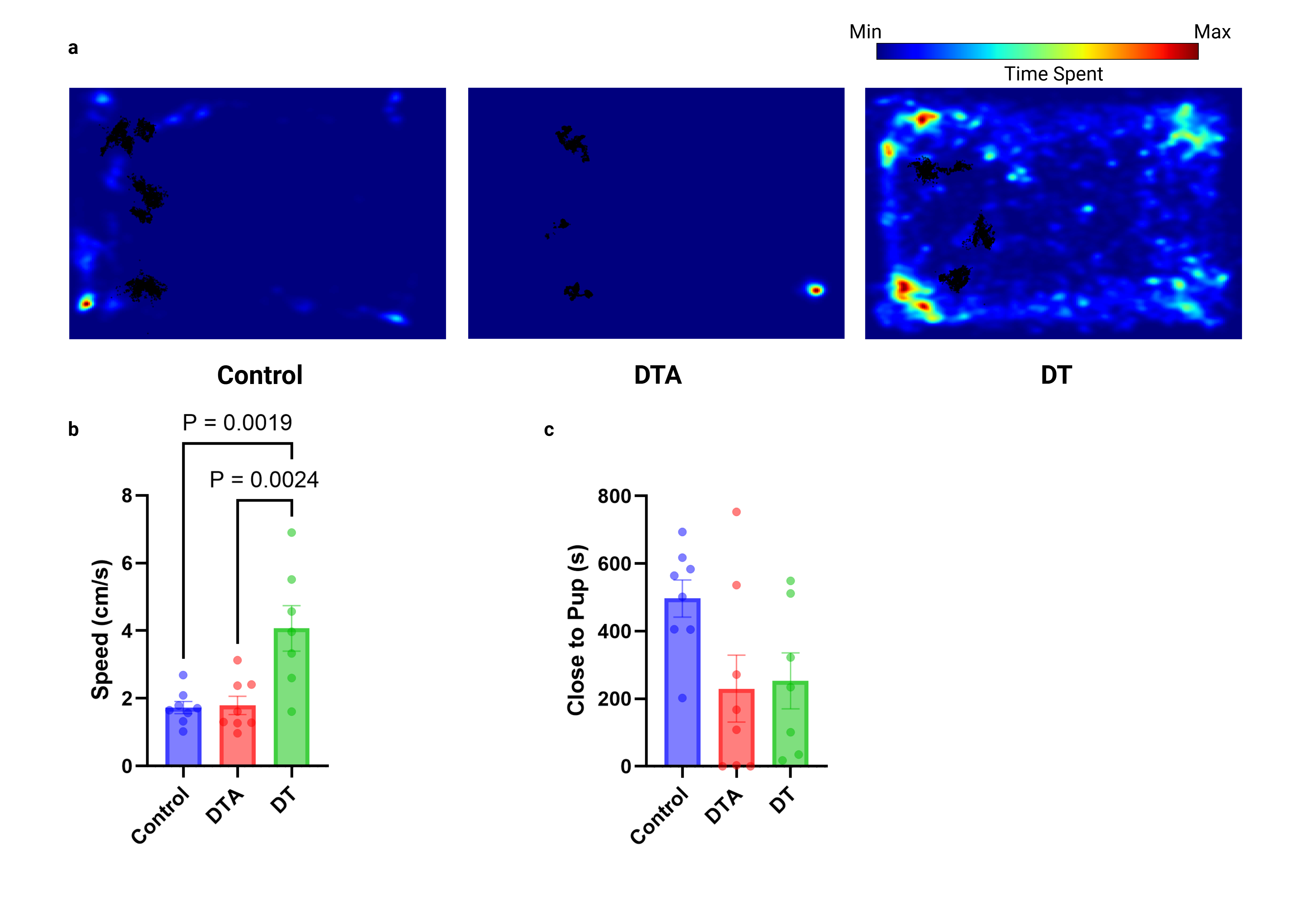

### Supplementary 3

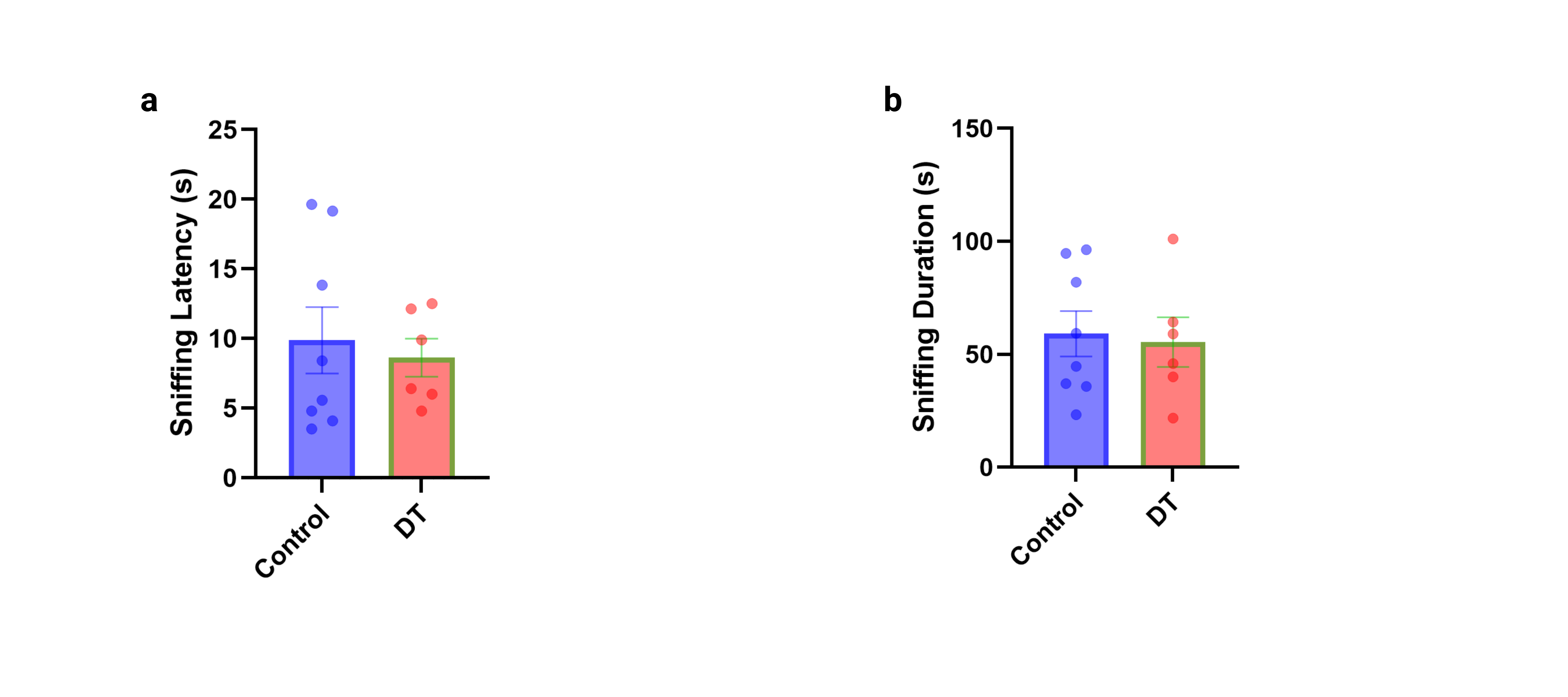

### Supplementary 5

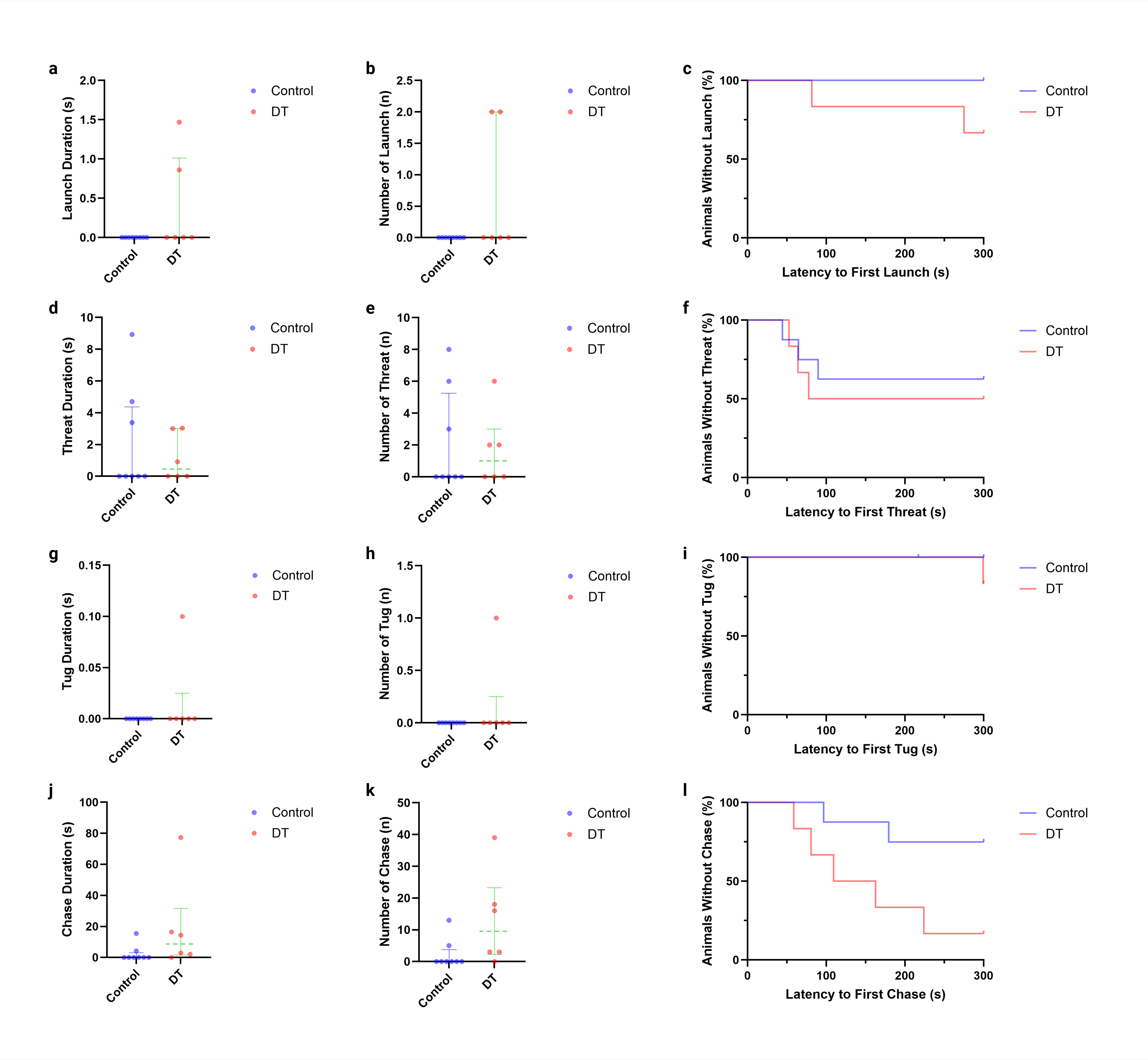
